# Observations on the interactions of mammals and birds with badger (*Meles meles*) dung pits at a site used by a single social group in an urban area

**DOI:** 10.1101/795484

**Authors:** Morgan Hughes, Scott Brown

## Abstract

During a monitoring study of a single social group of badger (*Meles meles*) at an urban site, incidental observations were noted of mammalian and avian species feeding within and removing material from *M. meles* dung pits. In response to these observations, infra-red cameras were deployed at dung pits for a 10-week period to document the nature, timing and frequency of these behaviours. Cameras were triggered a total of 954 times by a total of nine mammal and 12 bird species. Harvesting of material accounted for 28 % of latrine-associated behaviours. Results may have implications for disease transmission and the efficacy of badger surveys, particularly in areas where brown rats are prevalent.

## Introduction

The diet of the *M. meles* changes throughout the year as the species exhibits behavioural plasticity with regard to foraging, switching from a grain-based diet in summer to a largely fruit- and worm-based diet in autumn (Cheeseman & Neal 1998). *M. meles* digestive systems do not effectively process cellulose, suberin or lignin (Cheeseman & Neal 1998) and as such, undigested plant material can remain intact within faecal matter. Such material represents a readily available potential food source for foraging animals which is frequently renewed and deposited in a predictable place, as *M. meles* will re-use dung pits persistently (Roper, 2010). A single social group of *M. meles* at an urban Local Nature Reserve (LNR) in Walsall in the West Midlands was monitored using infra-red (IR) cameras by the authors from 2013 to 2018. Following incidental observations of mammalian and avian species apparently feeding within and removing material from dung pits, a 10-week study was undertaken to monitor activity around active dung pits to document the nature, timing, and frequency of the behaviour.

### The Site

The study site is a 13-hectare LNR situated in a fully urban context in Walsall in the West Midlands. The setts comprise a main sett (six entrances), an annex (two entrances), and several outliers and subsidiaries supporting a single social group of *M. meles* of approximately 13 individuals prior to annual dispersal (Hughes and Brown, 2017).

Classification of setts follows that of Kruuk (1978) and Thornton (1988). All setts experience regular anthropogenic disturbance including noise and vibration from nearby roads, industry, schools and sports grounds, walkers, dogs and cyclists. The main sett has been documented by the authors and local wildlife crime officers to have been subject to malicious disturbance including sett-blocking and attempted badger-baiting.

## Methods

### Settings

Five Bushnell HD Aggressor E2 Low-glow Trophycam Infra-red (IR) cameras were deployed facing either individual dung pits or latrine areas. Dung pits were selected based on known use by *M. meles* (i.e. those subject to deposition of faeces during the previous week) in areas away from public footpaths to reduce risk of human interference or camera theft. Cameras were deployed with no overlapping fields of view, set to record on PIR trigger, at high sensitivity, recording 20-second videos at 720p, with a 3-second delay. Batteries were changed when there was 1/3 battery power left or less. The 16 GB memory cards were formatted in-unit to improve write speeds (Wearn & Glover-Kapfer 2017). Cameras were checked weekly as per guidelines of a two-week checking schedule with an increased frequency in areas of high human activity (Ancrenaz et al. 2012). The survey comprised a total of 296 trap nights (Table 1) separated into 10 weekly segments commencing on 27/08/2017, corresponding to ISO week 35 (International Organization for Standardisation, 2017)). Each trap ‘day’ began at 00:00:00 hrs and ended at 23:59:59 hrs.

**Table 1:**
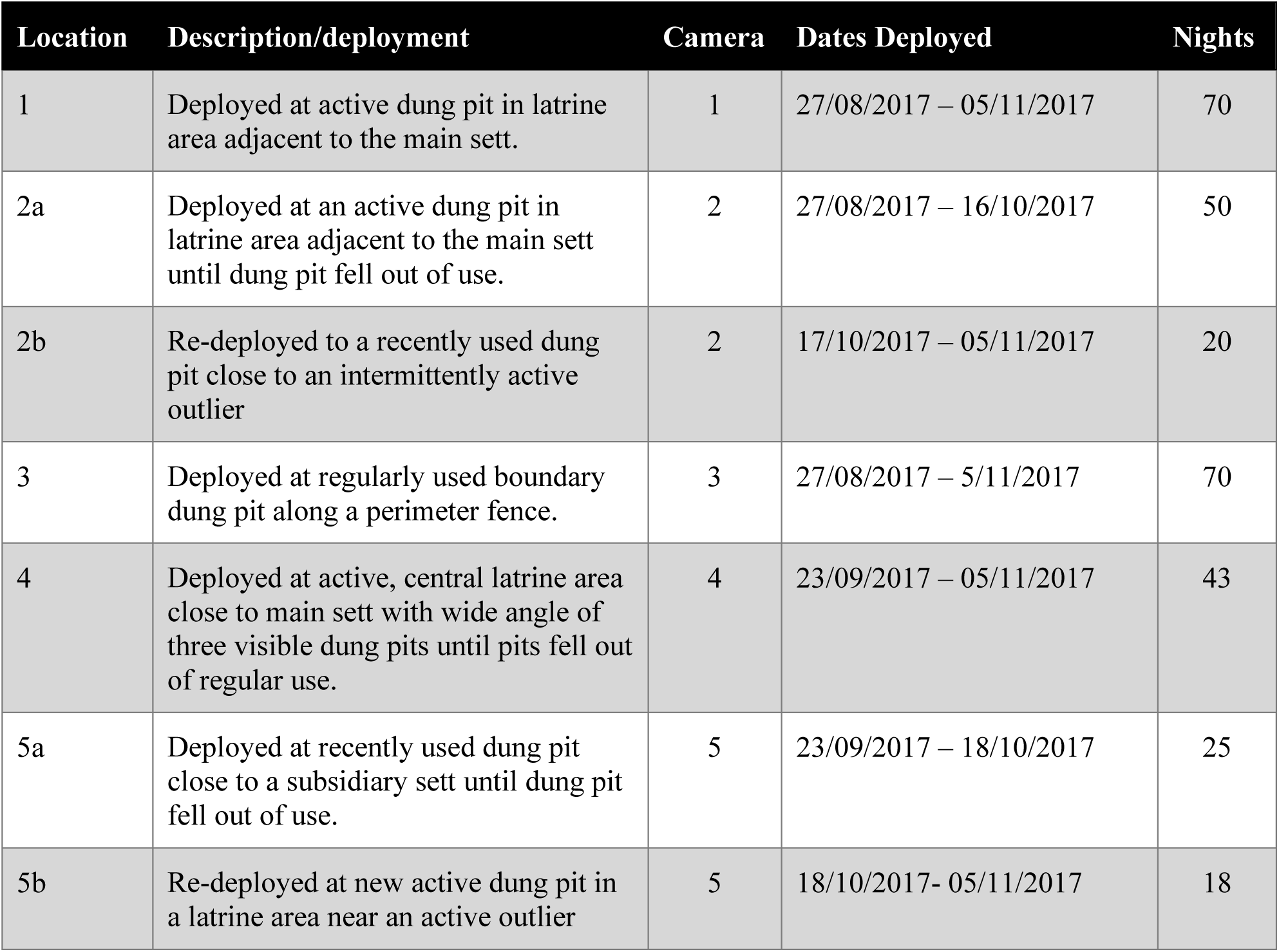
Camera deployment & trap nights.

| Location | Description/deployment | Camera | Dates Deployed | Nights |
| --- | --- | --- | --- | --- |
| 1 | Deployed at active dung pit in latrine area adjacent to the main sett. | 1 | 27/08/2017 – 05/11/2017 | 70 |
| 2a | Deployed at an active dung pit in latrine area adjacent to the main sett until dung pit fell out of use. | 2 | 27/08/2017 – 16/10/2017 | 50 |
| 2b | Re-deployed to a recently used dung pit close to an intermittently active outlier | 2 | 17/10/2017 – 05/11/2017 | 20 |
| 3 | Deployed at regularly used boundary dung pit along a perimeter fence. | 3 | 27/08/2017 – 5/11/2017 | 70 |
| 4 | Deployed at active, central latrine area close to main sett with wide angle of three visible dung pits until pits fell out of regular use. | 4 | 23/09/2017 – 05/11/2017 | 43 |
| 5a | Deployed at recently used dung pit close to a subsidiary sett until dung pit fell out of use. | 5 | 23/09/2017 – 18/10/2017 | 25 |
| 5b | Re-deployed at new active dung pit in a latrine area near an active outlier | 5 | 18/10/2017- 05/11/2017 | 18 |

### Data Management

For each time the cameras were triggered by animal movement, recording a video file (hereafter referred to as a ‘trigger’) the date, time, species, number of individuals and behaviour were recorded. Behaviours were classified (see **Table 2**) as either non-Latrine-Associated Behaviour (nLABs) comprising Commuting, Foraging, Caching and Camera Interaction or Latrine-Associated Behaviour (LABs) comprising Investigating, Toileting, Scent-marking and Harvesting.

**Table 2:** Ethogram of Behaviour Categories and Definitions.

| Behaviour Group | Behaviour Category | Definition |
| --- | --- | --- |
| **nLABs**<br>(non-Latrine-Associated Behaviours) | Commuting | uninterrupted movement through field of view of camera, showing no interaction with dung pits, immediate environment or camera |
|  | Foraging | behaviour associated with food-seeking |
|  | Caching | burying of food items |
|  | Camera Interaction | direct interaction with camera trap |
| **LABs**<br>(Latrine-Associated Behaviours) | Investigating | interaction with a dung pit not seen to be associated with Toileting, squat-marking, eating or stealing from it |
|  | Harvesting | foraging within a dung pit, eating within pit and/or carrying away faeces |
|  | Toileting | urinating or defecating on or in a dung pit |
|  | Scent Marking | squat-marking or spraying for purposes of olfactory communication |

It was not always possible to determine if an animal entering a dung pit was Harvesting due to the sightline of the camera or the direction of the animal’s egress. In the absence of direct evidence of Harvesting, such events were categorised as ‘Investigating’.

### Analysis

The statistical significance in differences in activity levels at the main sett between baseline and post-Toileting periods was calculated, using a χ2 test in SPSS (IBM Corp., 2016) based on trigger numbers over 21 consecutive days at the main sett.

## Results

Cameras were triggered by animals 954 times during the study, by a total of nine mammal species and 12 bird species. nLABs accounted for 78 % of triggers and LABs for 22 % of triggers (**Table 3**). nLABs comprised triggers of 45 % Commuting, 27 % Foraging, 4 % Caching and 1 % Camera Interaction. Only LABs were examined in detail. LABs comprised 14 % Investigating, 6 % Harvesting, 2 % Toileting and 1 % Scent-marking behaviours.

**Table 3:** Breakdown of Trigger Categorisation.

| Behaviour | No of Triggers | % of Triggers | % of Group Triggers |
| --- | --- | --- | --- |
| *Commuting* | 432 | 45% | 58 % of nLAB |
| *Foraging* | 257 | 27% | 35 % of nLAB |
| *Caching* | 42 | 4% | 6 % of nLAB |
| *Camera Interaction* | 11 | 1% | 2 % of nLAB |
| *Investigating* | 132 | 14% | 62 % of LAB |
| *Harvesting* | 60 | 6% | 28 % of LAB |
| *Toileting* | 14 | 2% | 7 % of LAB |
| *Scent Marking* | 6 | 0.6% | 2.8 % of LAB |

### Latrine Associated Behaviouors

Toileting occurrences were recorded 17 times, with 11 of those attributed to *M. meles* and the remainder being red fox (*Vulpes vulpes*), domestic cat (*Felix silvestris*) and dunnock (*Prunella modularis*) with four, one and one occurrences, respectively). Harvesting of material accounted for 28 % of LAB triggers. Of those, 82 % were by mammals comprising 77 % by brown rat (*Rattus norvegicus*), 3 % by grey squirrel (*Sciurus carolinensis*) and 2 % by wood mouse (*Apodemus sylvaticus*). Avian species accounted for 18 % of Harvesting triggers, comprising 13 % by magpie (*Pica pica*), and 2 % each by chaffinch (*Fringilla coelebs*), dunnock and wren (*Troglodytes troglodytes*).

There was a variability in Harvesting behaviour (**Figure 1**) with Harvesting representing 21 %, 30 % and 17 % of triggers in weeks 35, 37 and 38, respectively (with zero Harvesting activity being observed in week 36 due to camera failure). Lower levels of Harvesting (less than 5 % of triggers) took place during weeks 39 through 42, increasing again in weeks 43 and 44 to 6 % and 7 %, respectively.

**Figure 1:**
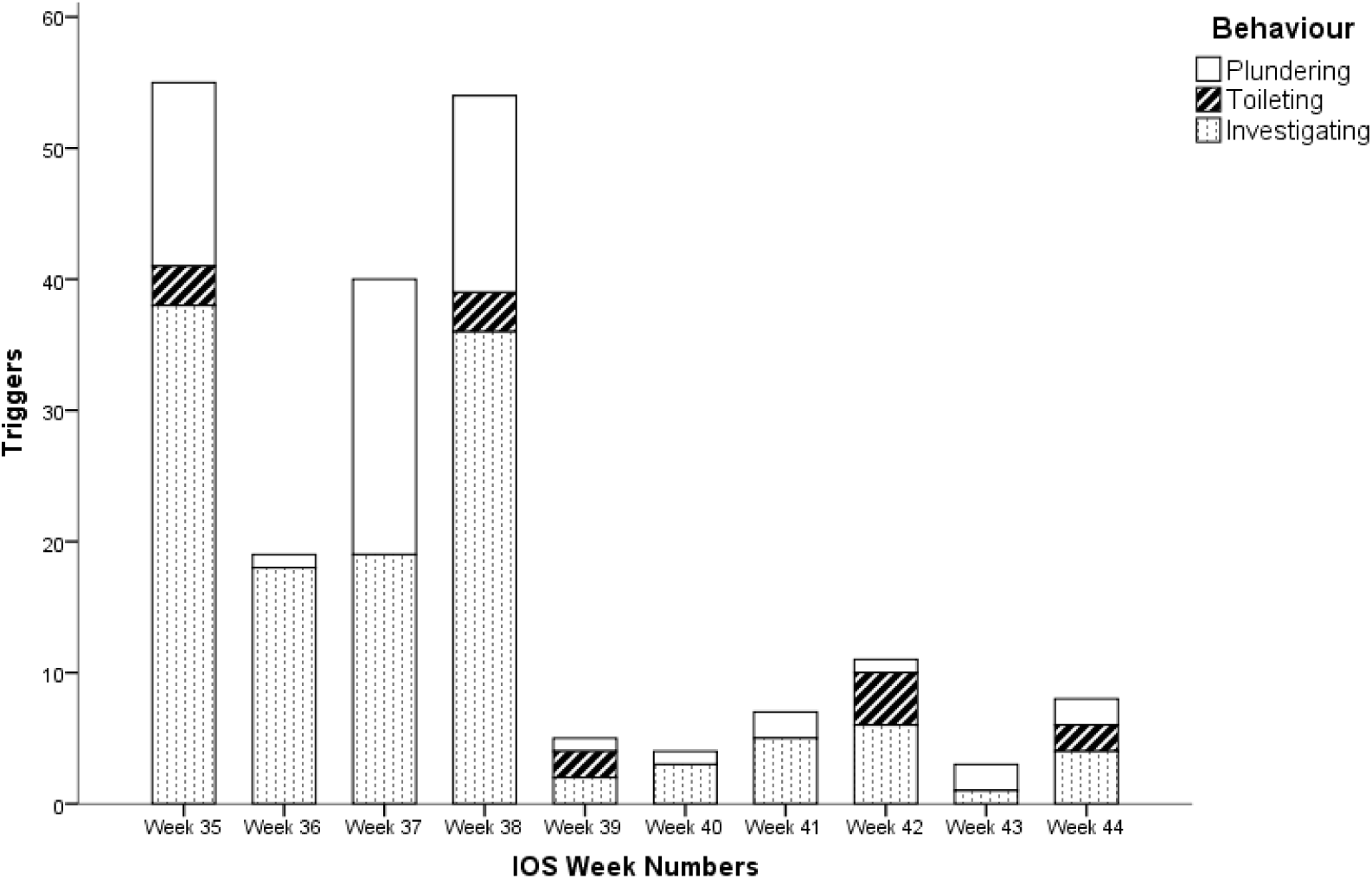
Triggers by Behaviour per week of the year (IOS, 2017)

### Post-Toileting Activity

Activity levels (all behaviours by all species) underwent an increase in the 24-hour period following Toileting events (**Figure 2**), with subsequent reduction in activity in the following 24-hour period down to a baseline of below 10 triggers per day over a 21-day period at the main sett in which four Toileting events were documented. This increase from baseline to post-Toileting activity (**Figure 3**, Figure 4) was subject to χ2 analysis indicating that this is unlikely to have occurred by chance (*p* <0.001).

**Figure 2:**
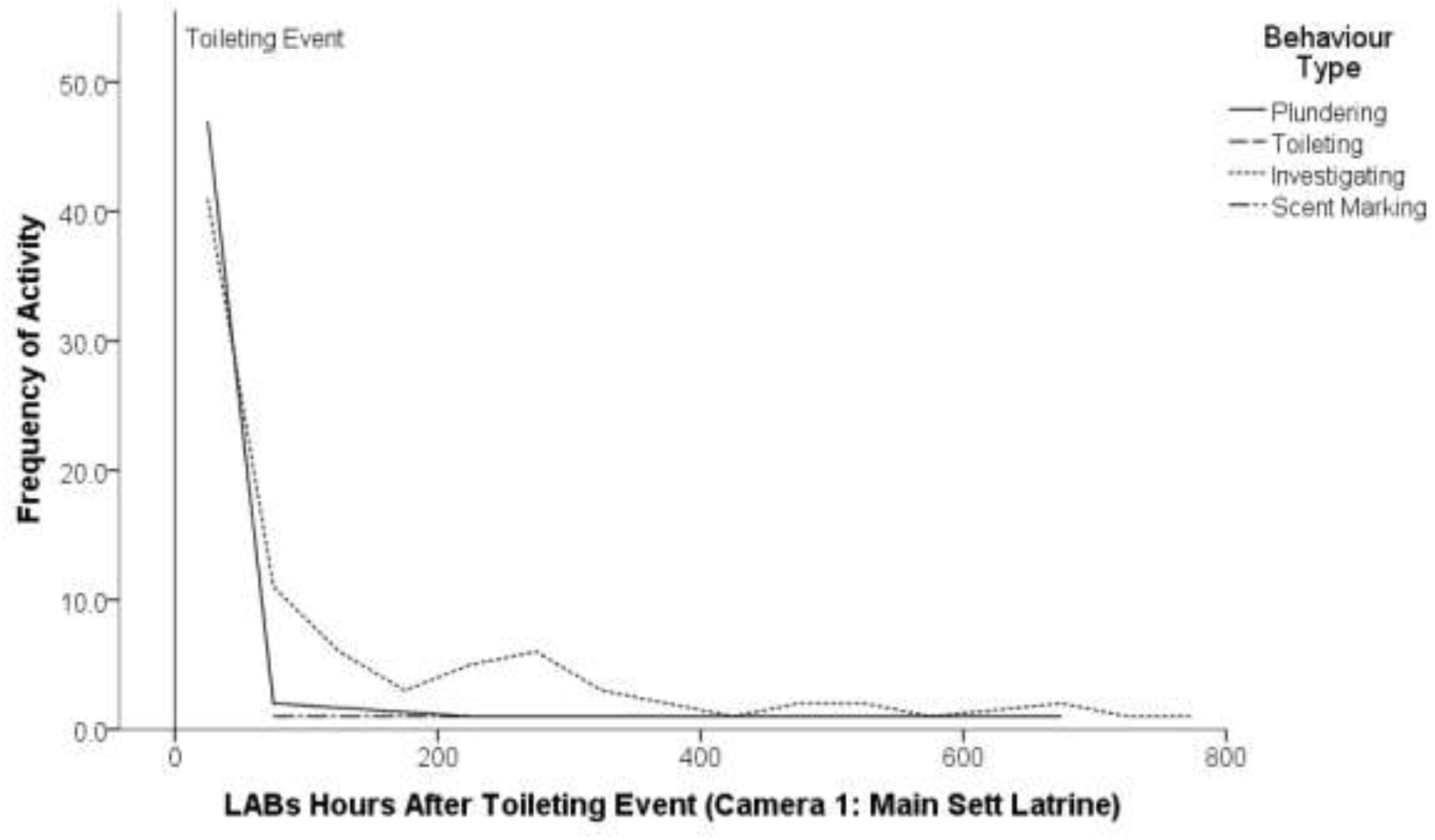
nLABs activity levels in hours after Toileting event/faecal deposit (triggers by all species at main sett, camera 1)

**Figure 3:**
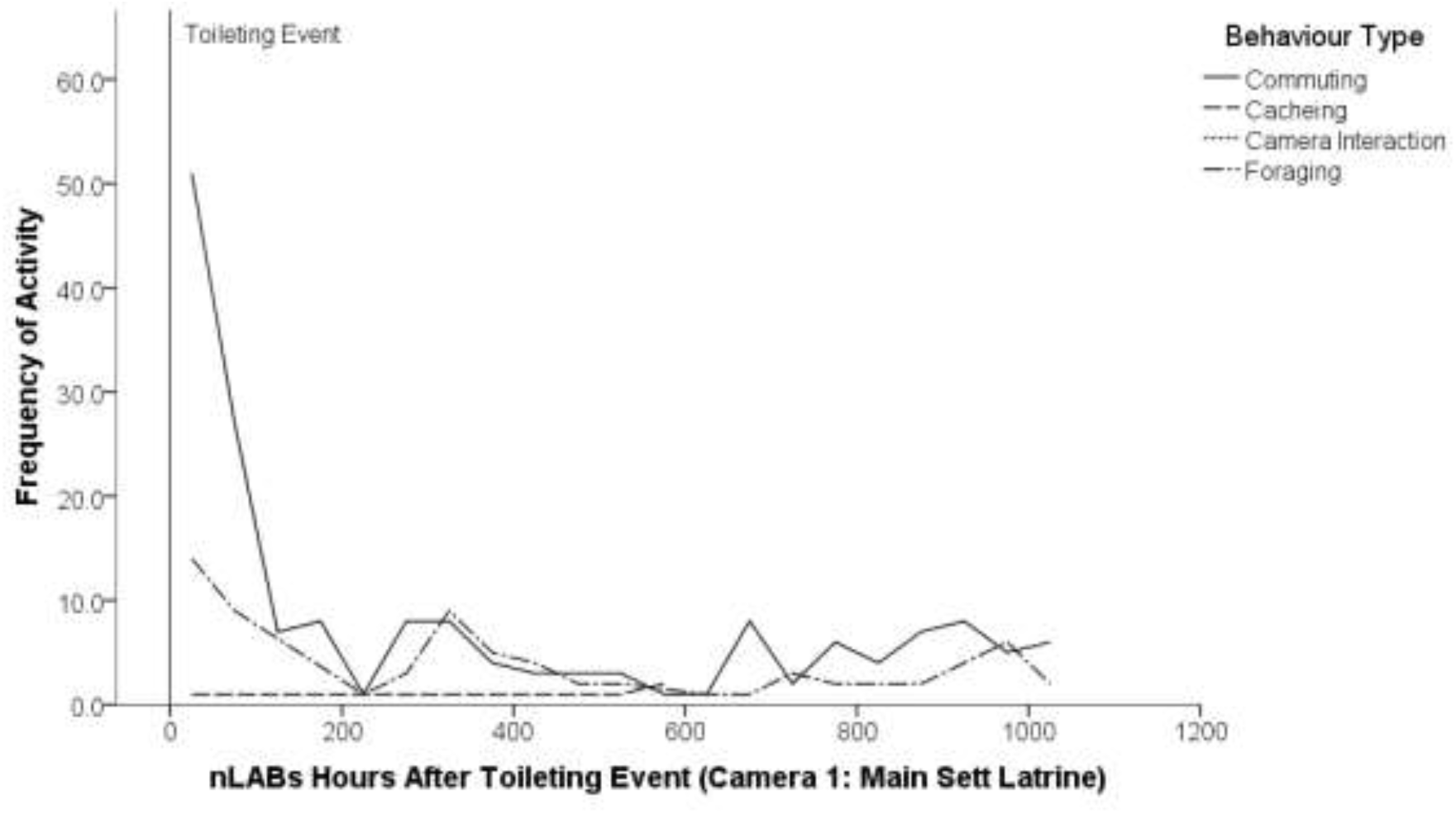
nLABs activity levels in hours after Toileting event/faecal deposit (triggers by all species at main sett, camera 1)

**Figure 4:**
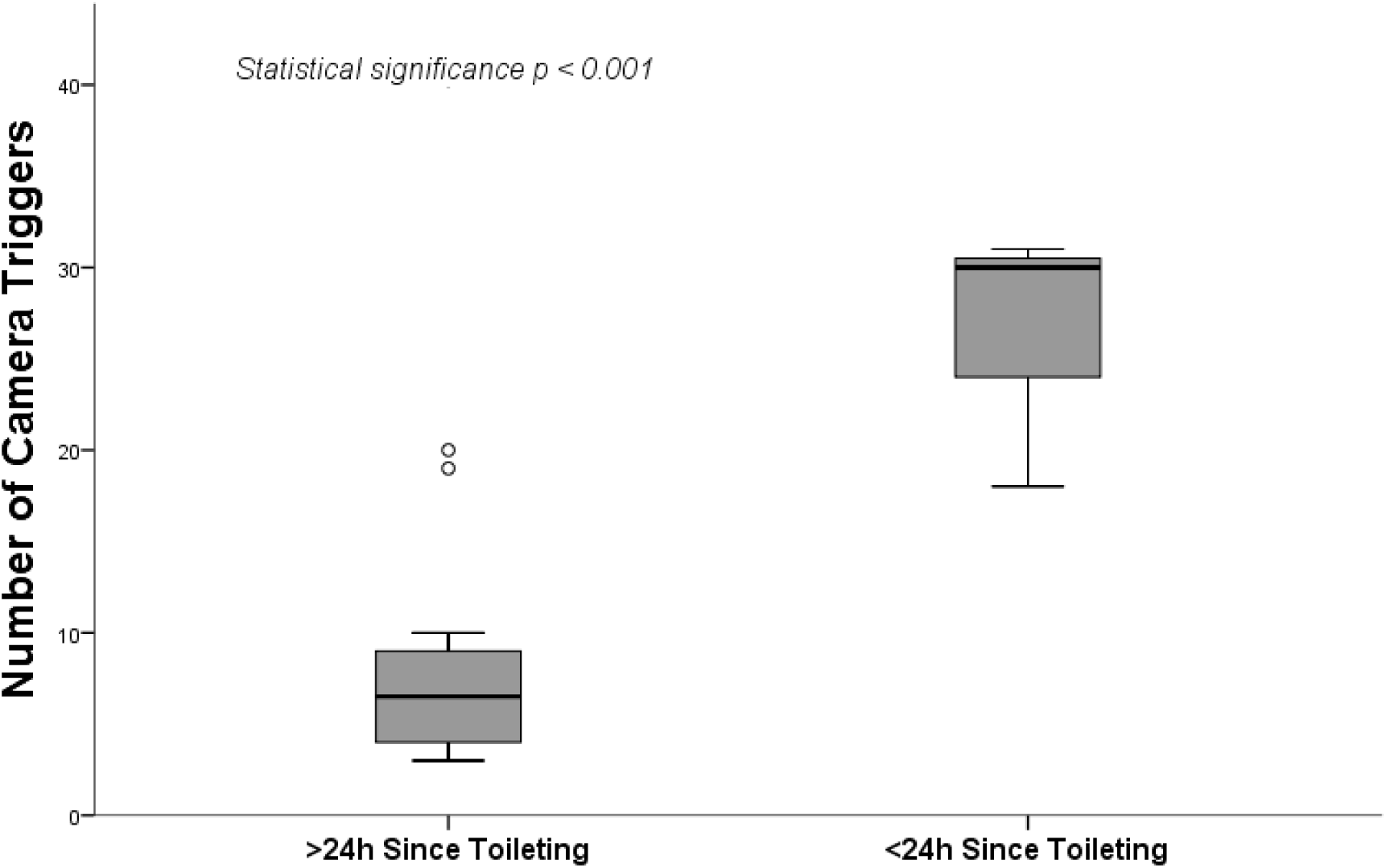
Box plots showing difference in activity levels at baseline (on days with >24 hours since faecal deposit) compared with days <24 hours since faecal deposit.

### Toileting vs. Harvesting

The majority (80 %) of *M. meles* Toileting activity took place between 03:00 and 05:30; the majority (85 %) of Harvesting activity by *R. norvegicus* took place between 08:00 and 09:00. On average, Harvesting commenced within two to six hours after faecal deposit and persisted until up to nine hours after deposit, with the majority of Harvesting happening in ‘events’ with multiple trips taking place to plunder a dung pit for faeces until the food resource was depleted, with subsequent investigatory trips to the dung pits. During one such event, 28 separate trips were recorded to a single dung pit by (presumed to be the same) adult male *R. norvegicus*. This is considered likely to have emptied the latrine of all solid faeces (**Plate 1, Plate 2**).

**Plate 1:**
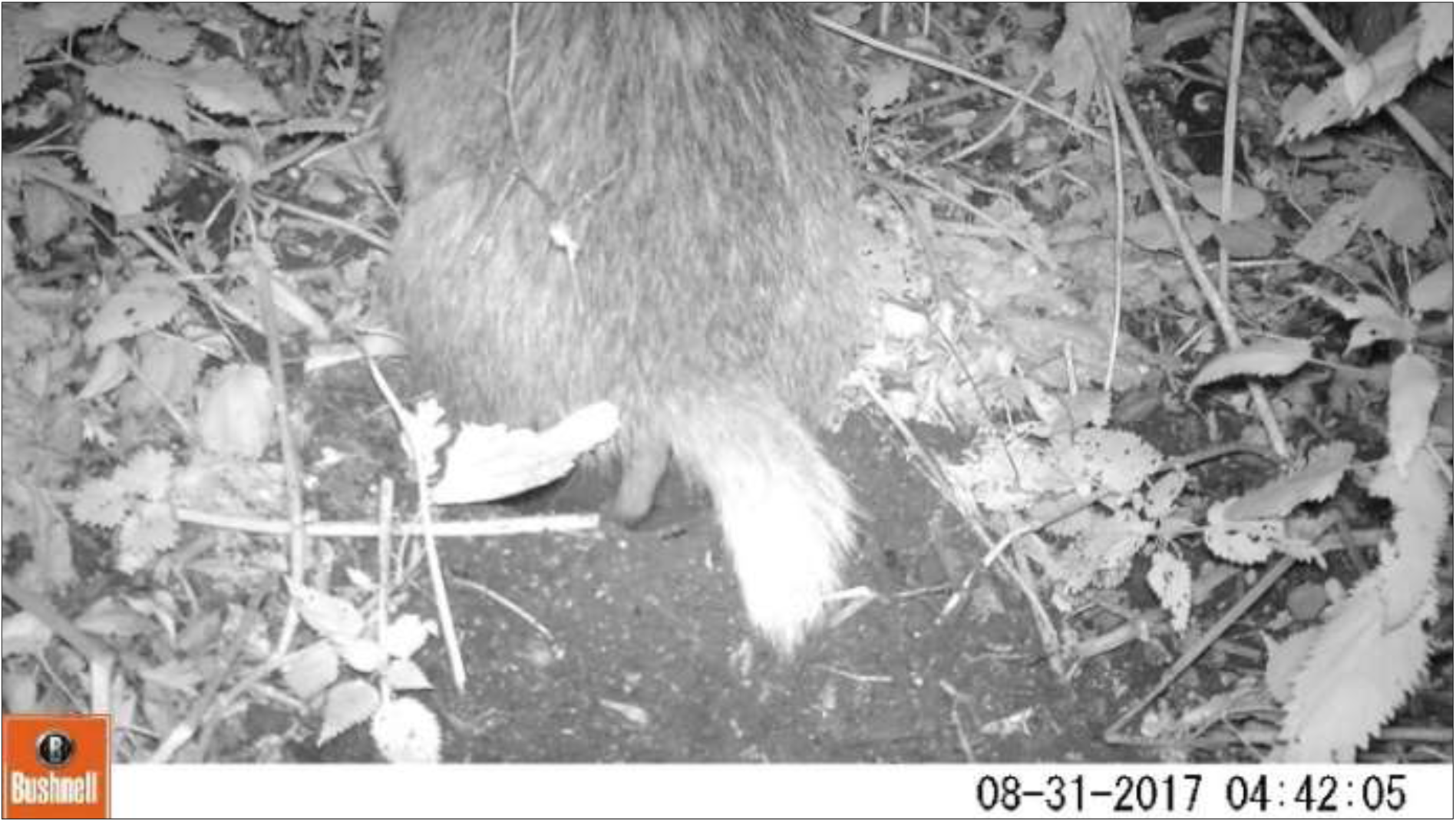
<u>M. meles</u> defecating into a dung pit at 04:42 © M Hughes & S. Brown 2017.

**Plate 2:**
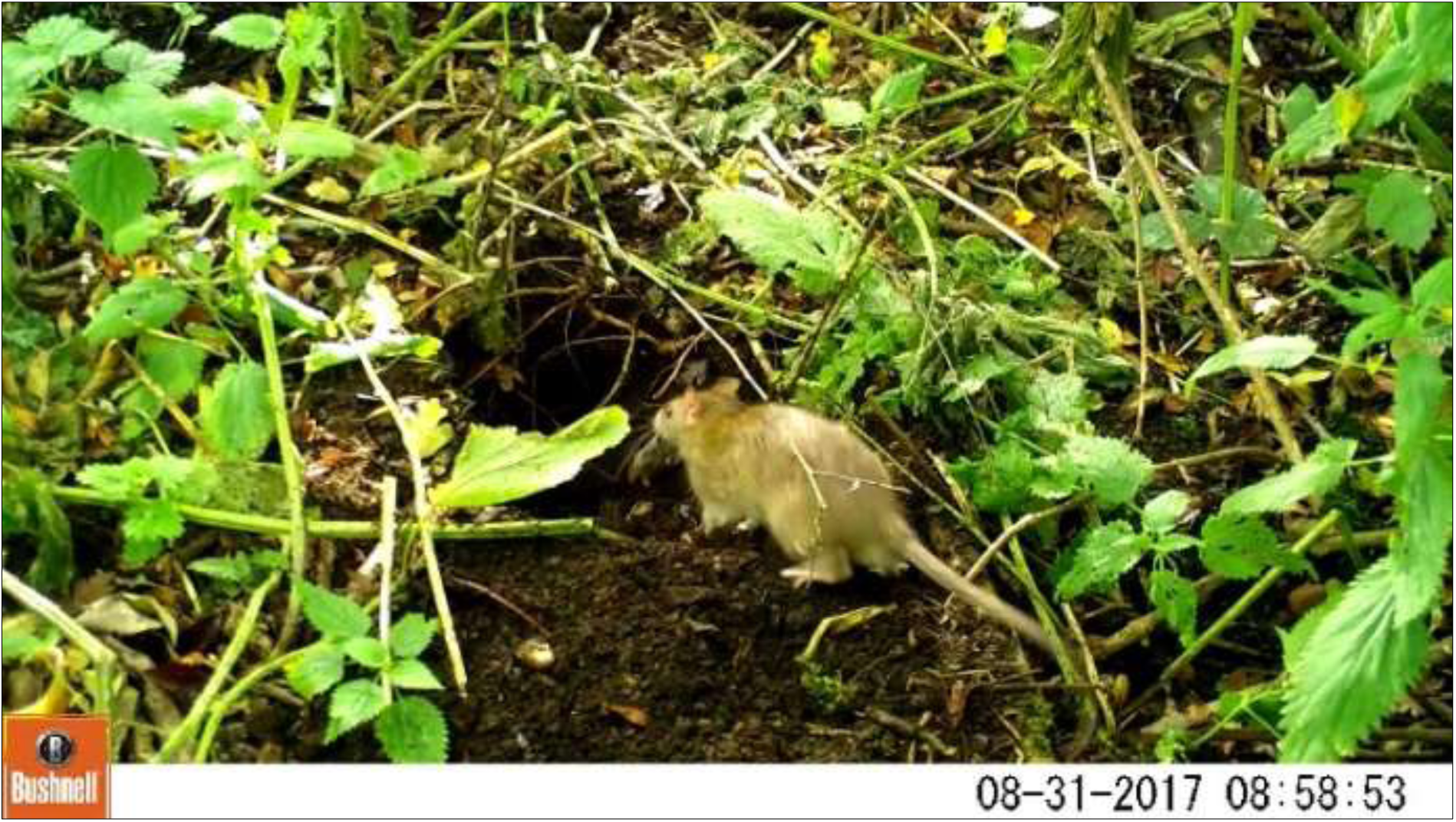
<u>R. norvegicus</u> Harvesting faeces at same dung pit 08:58 © M Hughes & S Brown 2017.

## Discussion

### Seasonality

Harvesting events were more frequent in the early part of the study (ISO weeks 35, 37 and 38). This may be due to seasonal *M. meles* dietary changes affecting faecal contents from those typical of grain-based diet in late summer **(Plate 3**) to those typical of a fruit and worm-based diet in early autumn (Cheeseman & Neal 1998). This dietary change would reduce the abundance of grains within faeces, making the dung pits a less lucrative food source.

**Plate 3:**
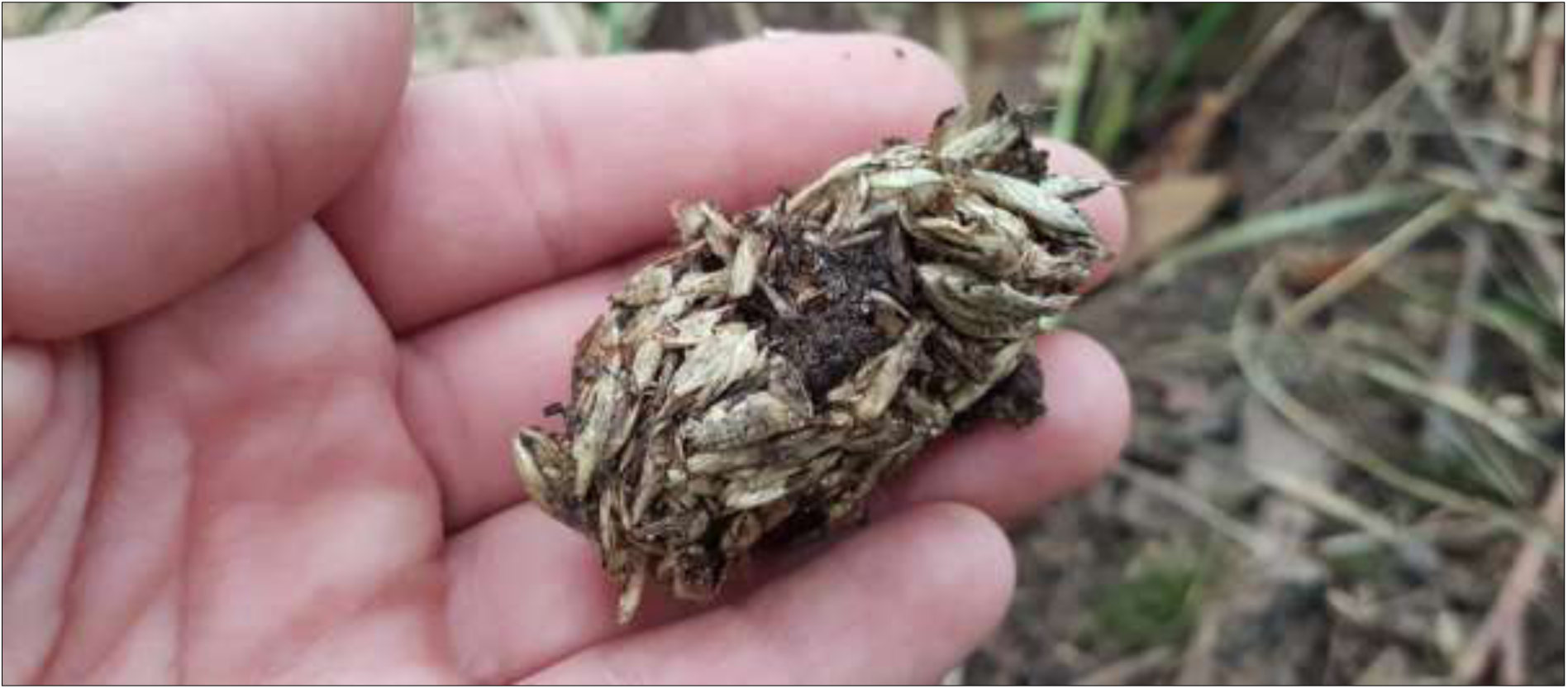
<u>M. meles</u> faeces with high grain content © Morgan Hughes 2017.

### Habitats

The observations noted during this study may be more prevalent urban environments (particularly in areas where feeding of wildlife takes place, such as in this study site) and arable environments (where the growth, harvesting and storage of grain crops take place) which are more likely to support larger populations of rodents than grasslands or woodlands where there is no supplementary artificial food source. Availability of supplementary food sources may also indirectly affect the prevalence of Harvesting behaviour due to its influence on the variability of *M. meles* diet.

### Disease Transmission

Should the behaviour documented in this study prove to be widespread, there may be implications to consider regarding disease transmission. For example, bovine tuberculosis (*Mycobacterium bovis*) has been documented to be present in both *R. norvegicus* and *A. sylvaticus* (Little et al., 1982; Delahay et al., 2001); *R. norvegicus* is known to carry other diseases that are transmissible to cattle (Ward et al., 2006). While it has been acknowledged that *R. norvegicus* is a potential vector for transmission of *M. bovis* in agricultural landscapes due to the frequency of their contact with livestock and contaminated food stores (Delahay et al., 2001), their potential to transmit diseases between social groups of *M. meles* in urban environments is little understood.

### Survey Efficacy

Presence of faeces in dung pits is typically used as an indication of *M. meles* presence and activity, as well as social group size (Wilson et al., 1997). Current methods for analyzing *M. meles* territories rely on bait marking (Delahay et al., 2000). The findings of this study indicate that in areas where *R. norvegicus* populations are more prevalent, the survey efficacy of *M. meles* activity (Reynolds & Harris 2005) and bait marking surveys (Delahay et al. 2000) may be adversely affected by the Harvesting behaviour exhibited by *R. norvegicus* as described here, particularly in incidences where entire faeces are removed or dung pits are emptied. Current protocols suggest placing bait in late afternoon to reduce the consumption of bait by diurnal, non-target species, but there are no times stipulated for checking of latrines (Delahay et al. 2000), which is typically undertaken at the same time as the visit to place bait. The results of this study indicate that survey efficacy may be improved by undertaking checks of latrines as early as possible in the day in order to maximise the chance of finding faecal matter in dung pits, particularly at times of year when *M. meles* diet is grain-based.

Current bait marking methodology (Delahay et al. 2000) suggests an optimal survey period of February - April, and a second survey period in September-October. Harvesting behaviour is more likely to take place during times of the year when faeces contain grains and are more solid. As such, is likely to be less of a constraint in October surveys. However, the provision of bait itself may trigger an increase in harvesting activity and make faeces more viscous and able to be removed.

### Limitations

Equipment failure of unknown causes occurred on two occasions, resulting in loss of data. Intermittent failure of cameras to trigger has also been observed, which may contribute to the number of false negatives. When triggered, IR cameras used produce an audible click, which has been demonstrated by to be detectable by mammals (Meek et al. 2014), possibly causing mammals to alter their normal behaviour. Some Mustelidae are able to detect light with IR wavelengths of up to ∼870 nm (Newbold & King 2009). The cameras used for this study use IR light of 850 nm. Individual *M. meles* have been observed by the authors to turn towards cameras when triggered. It is unknown whether the animals’ response is to the light, the click, or to both (Meek et al. 2016). Ancrenaz et al. (2012) report that most high-end, passive IR sensors can detect animals as small as 100 grams within 2 m of the sensor. Trigger success rates of high-end cameras (Glen et al. 2013) using 1080p no-glow cameras with trigger speeds of 0.2–2.1 seconds to detect stoats (*Mustela erminea*) were up to 80 % successful, depending on the animal’s speed. The stoat weighs 140 – 445 g, and is an appropriate analogue for *R. norvegicus* at 200-300 g (Mammal Society 2017).

### Recommendations

A constraint to the study is the small sample size, which took place at a single site used by a single social group. Further study is required to ascertain whether this behaviour can be readily observed in other urban areas, agricultural areas and in non-anthropogenic habitats.

